# Contrasting genomic trajectories of *Bartonellaceae* symbionts of planthoppers

**DOI:** 10.1101/2025.08.24.672000

**Authors:** Mingjie Ma, Anna Michalik, Junchen Deng, Yi Hu, Piotr Łukasik

## Abstract

Symbioses with microorganisms have shaped the nutritional biology and evolution of many insects. For example, several ant clades have adapted to nutrient-poor diets through symbiosis with a specific clade of bacteria in the family *Bartonellaceae* (Hyphomicrobiales), notorious for also including virulent vertebrate pathogens. Here we show that *Bartonellaceae* phylogenetically placed within the clade that has only encompassed ant symbionts to date – *Candidatus* genus Tokpelaia – have established as symbionts in four different clades of planthoppers (Insecta: Hemiptera: Fulgoromorpha). Genome size and contents indicate different levels of integration of these strains into the planthopper host biology and their diverse roles. Symbionts infecting one of the clades have some of the largest genomes among *Bartonellaceae*, at ca. 2 Mb, two others are under 700 kb, and the fourth is reduced to barely 158 kb. The planthopper-associated *Tokpelaia* strains with larger genomes, similarly to ant symbionts, encode multiple amino acid and vitamin biosynthesis genes, complementing the degraded nutritional capabilities of their hosts’ ancient heritable endosymbionts. Strikingly, the smallest *Tokpelaia* genome lacks any genes linked to essential amino acid biosynthesis, in contrast to all other known insect-associated bacteria with genomes of comparable size. We identified a single vitamin biosynthesis gene and iron-sulfur cluster assembly genes as its only putative contributions to the host biology.

Our results broaden the host spectrum of non-pathogenic *Bartonellaceae*, indicating that they have contributed to nutrition and symbiotic consortium function in diverse diet-restricted host clades. They also highlight an unexpectedly broad range of evolutionary outcomes for this important bacterial group.

**Significance:** Many insects rely on bacterial partners to overcome nutrient-poor diets, yet the diversity and evolutionary trajectories and outcomes for these associations remain unclear. Our discovery of *Bartonellaceae* bacteria in planthoppers improves the understanding of the host range of these broadly relevant bacteria best-known as vertebrate pathogens. The finding that these planthopper symbionts generally have the capacity to produce essential amino acids and vitamins relevant to hosts and likely form long-term symbiotic associations agree with the expectations for sap-sucking hemipteran symbionts. However, the breadth of sizes and functions of the newly assembled genomes substantially expand the known range of states for *Bartonellaceae*, and their evolutionary trajectories. In particular, the identification of a strain with an ultra-reduced genome of only 158kb with no amino acid biosynthetic functions contrasts with all other known insect-symbiotic bacteria with genomes in a similar size range. This study also indicates how symbionts may enable the degeneration, functional loss, and evolution of other bacteria residing in the same host.

## 1. Introduction

Microbial symbionts can play dramatically important and highly diverse roles in the biology of their animal hosts, shaping their life history traits, ecological niches, biological interactions, and evolutionary processes (Moran 2007; Douglas 2014; Lukasik and Kolasa 2024). In particular, host long-term evolutionary trajectories have repeatedly been shaped by the microbial contributions to the host nutrition (Moran 2007; Sudakaran, et al. 2017; Brentassi and de la Fuente 2024). Microbial symbionts frequently help address challenges related to the digestion of recalcitrant foods, the neutralization of toxic compounds, and the acquisition of limiting nutrients that animals require but cannot make on their own. For example, plant-based foods typically have low nitrogen contents and specifically lack essential amino acids as well as vitamins, and many herbivores rely on symbiotic bacteria for nitrogen recycling, upgrading, and sometimes fixation, and for the biosynthesis of lacking nutrients (Hansen and Moran 2014). Insects provide many independent examples of associations between heritable symbiont acquisitions and shifts to limiting diets (Cornwallis, et al. 2023).

Ants (Hymenoptera: Formicidae) are often quoted as an example of a clade that has undergone extensive dietary diversification, including multiple independent transitions from more balanced omnivorous diets to herbivory (Russell, et al. 2009; Smith, et al. 2023). These nutritional changes have typically been enabled by bacterial symbionts, most often in the family *Bartonellaceae* (Alphaproteobacteria: Hyphomicrobiales) (Russell, et al. 2009; Bisch, et al. 2018). Genomics and experiments have confirmed *Bartonellaceae* roles in herbivorous genera *Cephalotes* (Hu, et al. 2018), *Dolichoderus* (Bisch, et al. 2018), and *Tetraponera* (Ma, et al. 2025), where they help recycle nitrogenous waste and supplement the host with nitrogen-rich compounds, especially amino acids. On the other hand, *Bartonellaceae* were also found in carnivores, and in the predatory ant *Harpegnathos saltator* they encode genes related to protein degradation, amino acid transport, and riboflavin biosynthesis, apparently assisting the hosts with the digestion of protein-rich diets and using amino acids as energy source, in addition to provisioning vitamins (Neuvonen, et al. 2016; Bisch, et al. 2018). Phylogenomic analyses have shown that the functionally diverse ant symbionts form a monophyletic clade – *Candidatus* genus *Tokpelaia* – that is sister to the genus *Bartonella* that contains notorious human and mammalian pathogens (Segers, et al. 2017; Wagner and Dehio 2019), but also symbionts of bees and other arthropods (Bisch, et al. 2018; Ma, et al. 2025).

Planthoppers (Hemiptera: Fulgoromorpha) showcase a very different nutritional history and symbiotic make-up. They have ancestrally fed on phloem sap, and to counteract its imbalanced nutritional contents, have repeatedly formed symbiotic associations with microorganisms that provide the lacking nutrients (Bennett and Moran 2013; Bell-Roberts, et al. 2019; Brentassi and de la Fuente 2024). Specifically, this 263-million-year-old infraorder (Deng, et al. 2025) has ancestrally associated with a Bacteroidetes symbiont *Candidatus* Sulcia muelleri (further referred to as *Sulcia*) and a Betaproteobacteria symbiont *Ca.* Vidania fulgoroidea (thereafter: *Vidania*) (Urban and Cryan 2012; Deng, et al. 2023). These two symbionts, transmitted strictly maternally and co-diversifying with their planthopper hosts (Moran, et al. 2005), jointly synthesize the 10 essential amino acids. In most planthoppers, *Sulcia* produces 3 [leucine, valine and isoleucine], and *Vidania* 7 [methionine, histidine, tryptophan, threonine, lysine, arginine and phenylalanine] (Michalik, et al. 2021; Michalik, et al. 2025). However, in several planthopper clades, these microbes have been replaced by Hypocreales fungi (Siehl, et al. 2024; Michalik, et al. 2025), or else, joined by additional microbes that established residence within host tissues alongside the ancient endosymbionts (Michalik, et al. 2023; Brentassi and de la Fuente 2024). These additional symbionts, representing the genera *Sodalis*, *Asenophonus*, *Serratia*, and others, can complement the amino acid biosynthesis pathways of the ancestral symbionts and frequently encode genes involved in the biosynthesis of B vitamins (Bennett and Mao 2018; Michalik, et al. 2021; Michalik, et al. 2024). However, we do not have a good understanding of how the nutritional functions vary among symbiont or planthopper clades, how they have evolved, and how they complement the functions encoded by other members of the symbiotic partnership. As the genomics-based characterization of these more recently acquired symbionts in Auchenorrhyncha has been largely limited to a few species carrying *Sodalis* (Koga, et al. 2013; McCutcheon, et al. 2019; Michalik, et al. 2021), we know little about other microbes forming relationships with these sap feeders.

In our recent metagenomics study spanning 149 species of taxonomically diverse planthoppers from different geographic areas (Deng, et al. 2025; Michalik, et al. 2025), in some of the surveyed species we identified, for the first time, *Bartonellaceae* symbionts. This included a strain infecting *Saccharosydne* sp., a Delphacidae planthopper, which had lost *Sulcia* and whose *Vidania* had the smallest genome of any known bacteria, barely 50 kilobases, reduced to provisioning a single amino acid, phenylalanine (Michalik, et al. 2025). We thus postulated that in this and other systems, the nutritional functions lost from the ancient symbionts’ genomes may have been taken over by the *Bartonellaceae*. This conclusion has prompted a closer investigation into the nature and significance of the associations between *Bartonellaceae,* their planthopper hosts, and co-residing other microbes.

Specifically, we asked about *Bartonellaceae* distribution across planthopper clades and the phylogenetic relationships among the newly identified strains and other members of the family, including mammalian pathogens and ant symbionts. We also asked about the genomic characteristics and contents of these newly identified strains, particularly in relation to nutritional provisioning that had been identified as a key function of previously studied *Tokpelaia* symbionts of ants. Finally, we asked how these microbes fit into multi-partner symbioses that provide hosts with essential nutrients. We addressed these questions by exploring the extensive recent metagenomics data for diverse global planthoppers, and through comparative genomics analyses of the newly identified planthopper-associated and previously published *Bartonellaceae* strains.

## 2. Results

### 2.1. The distribution and phylogenetic relationships of planthopper-associated *Bartonellaceae*

The systematic survey of 149 phylogenetically diverse planthopper species (Deng, et al. 2025; Michalik, et al. 2025) using 16S rRNA sequences reconstructed from metagenomes using phyloFlash (Gruber-Vodicka, et al. 2020), revealed *Bartonellaceae* in six species from four planthopper families, all classified to the genus level only. They included three members of the family Cixiidae: two representatives of the genus *Bothriocera* (BOTSP1, BOTSP2) and *Oecleus* sp. (OECLEU); a Dictyopharidae species *Lyncides* sp. (LYNSP); a Delphacidae planthopper *Saccharosydne* sp. (SACSP1), and a Kinnaridae species *Oeclidius* sp. (OECLID) (Fig. 1A, Tab. S1). This, combined with the phylogenetic distance among *Bartonellaceae*-hosting planthoppers (see below) indicated at least four independent host-microbe associations – one in each family. In turn, the Cixiidae clade that included three *Bartonellaceae*-hosting species also contained two species with no traces of this bacterium, and based on currently available data, we cannot distinguish among scenarios such as independent infections, an ancestral gain followed by subsequent losses in some clades, or a facultative association.

**Fig. 1.**
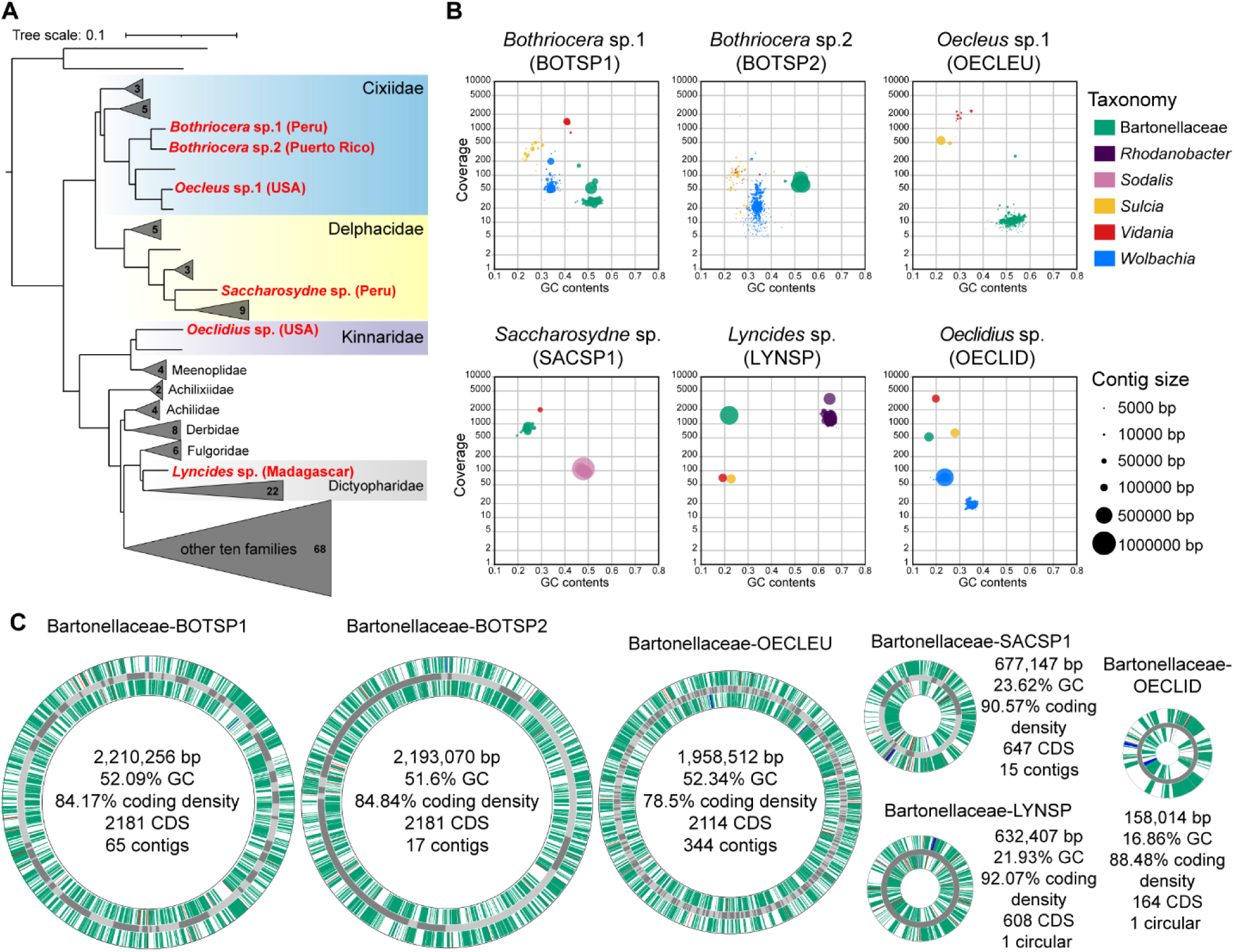
*Bartonellaceae* symbiont distribution across planthoppers, their representation within metagenomic assemblies, and genomic characteristics. **(A)** The genome-based phylogeny of 149 planthopper species representing 19 families, modified from Deng, et al. (2025), with species harboring *Bartonellaceae* indicated with red font. Clades where these symbionts were not detected were collapsed, with the number of species in these clades shown. **(B)** Taxon-annotated GC content-coverage plots for metagenomic assemblies of six planthopper species where *Bartonellaceae* were identified. Colored blobs represent contigs corresponding to the identified genomic fragments of different symbiotic bacteria. The size of each blob represents contig size. **(C)** Circular diagrams of *Tokpelaia* genomes from six species, showing gene positions on forward and reverse strands and basic genome characteristics. Note that only two genomes were circularized - most comprise multiple contigs, connected for the ease of illustration. The inner and outer circles represent coding sequences (CDS), and the functional categories are shown in green. tRNA, rRNA, and tmRNA genes are shown in red, blue, and purple, respectively. The dark gray and light gray in the middle circle represent different contigs. The size of the first five circles correlates with genome size, the sixth (OECLID) is magnified three times.

The metagenome assemblies for these six planthopper species contained a set of contigs, between 1 and 344, with significant sequence similarity to the genome of *Candidatus* Tokpelaia hoelldoblerii (*Bartonellaceae*) from a predatory ant *Harpegnathos saltator* (Neuvonen, et al. 2016), and forming a distinct cluster in GC contents and coverage space (Fig. 1B, Tab. S2, S3). The *Bartonellaceae* symbionts from Cixiidae planthoppers (BOTSP1, BOTSP2, and OECLEU) all possessed relatively large genomes, ranging from 1.96 to 2.21 Mb across 17-344 contigs, predicted to encode 2,114–2,181 proteins, and with the coding density from 78.5% to 84.84% (Fig. 1C). Their genomic GC content values were relatively high, between 51.6% and 52.34%. The *Bartonellaceae* symbionts of SACSP1 and LYNSP had genomes approximately three-fold smaller, 0.68 Mb (15 contigs - SACSP1) and 0.63 Mb (circular - LYNSP), with 647 and 608 predicted protein-coding genes, respectively. These reduced genomes also exhibited markedly higher coding densities—90.57% for SACSP1 and 93.2% for LYNSP—as well as much lower GC contents, 23.62% and 21.93%, respectively (Fig. 1C, Tab. S2). In turn, for the Kinnaridae strain OECLID, we have identified a single circularly mapping contig of 158 kb, with a GC content of 16.86% and a coding density of 88.48%, encoding predicted 164 genes that included the large majority of those conserved in other extremely reduced bacterial genomes (McCutcheon, et al. 2024). Despite the low sequence similarity to any previously known *Bartonellaceae* (84.13 - 86.39% at 16S rRNA gene, amino acid identity across phylogenetically informative genes: 47.03 - 51.47%, Tab. S4, S5) amino acid sequences of protein-coding genes primarily resembled *Bartonellaceae* among all NCBI records. No further contigs within the assembly could be classified to the same genome.

Phylogenomic analysis of the planthopper-associated *Bartonellaceae* reliably placed them among previously characterized ant symbionts forming the genus *Tokpelaia* (Fig. 2A, S1). Specifically, the planthopper strains form two well-supported subclades within *Tokpelaia*. One of them, on a relatively short branch, comprises symbionts of Cixiidae planthoppers (BOTSP1, BOTSP2, and OECLEU) and forms a sister clade with strains of *Tetraponera* ants. The second clade, monophyletic but with very long branches among strains, comprises symbionts of LYNSP, SACSP1, and OECLID. These small-genome strains show a closer relationship to smallest-genome symbionts of *Dolichoderus* ants, with strong support but long branches. This indicates an accelerated rate of sequence evolution, which combined with a strong AT bias, could have affected the accuracy of phylogenomic reconstructions through systematic problems such as long branch attraction, especially for the extremely reduced OECLID (Susko and Roger 2021). 16S rRNA gene-based phylogeny, including a much broader sampling of *Tokpelaia* sequences, produced a topology broadly consistent with the genome-based analysis, again placing the planthopper-associated symbionts among ant-associated *Tokpelaia* strains and forming two separate clades (Fig. 2B, S2).

**Fig. 2.**
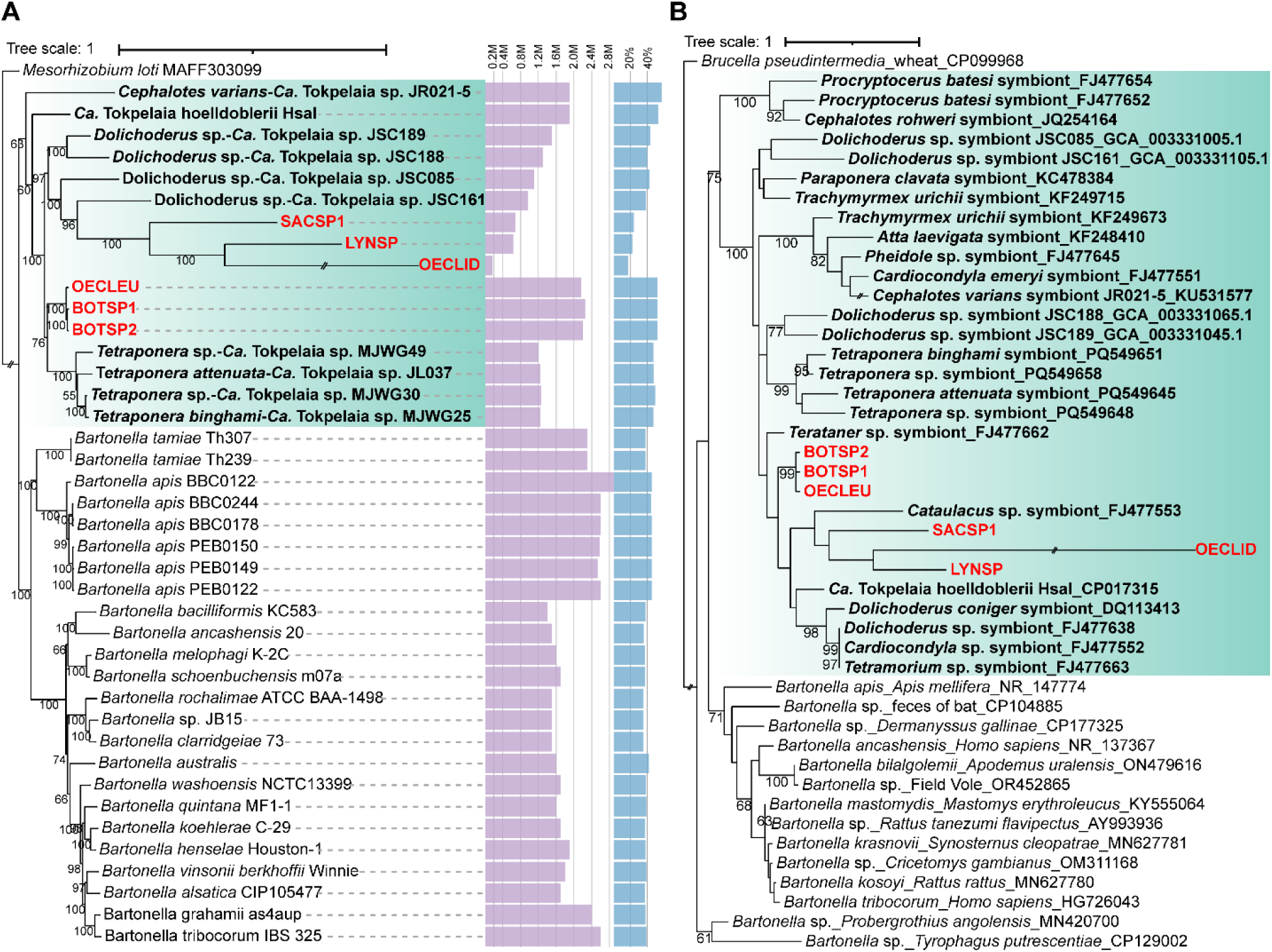
Phylogenetic placement of *Bartonellaceae* symbionts of planthoppers. **(A)** Maximum likelihood analysis of *Bartonellaceae* strains with sequenced genomes, based on 68 homologous single-copy genes. The purple bars represent the genome size, and the blue bars - the GC content. **(B)** Maximum likelihood phylogeny of full-length 16S rRNA gene sequences of *Bartonellaceae* associated with planthoppers, ants, and selected other hosts, derived from metagenomes and the NCBI database. In both panels, the *Tokpelaia* clade is indicated with green boxes; bold black labels indicate ant-associated, and bold red – planthopper-associated strains. Bootstrap support values above 50% are shown by the respective nodes. The very long branch leading to the OECLID strain was truncated in both figures.

The genome synteny comparisons did not support phylogenetics-based conclusions (Fig. S3). The highly fragmented assemblies of two of Cixiidae symbiont genomes prevented comparisons within that clade, and there was little similarity among strains comprising the second clade. Surprisingly, large syntenic regions were detected between BOTSP2 and SACSP1, despite them representing different clades. Also, symbionts from BOTSP1 and BOTSP2 shared limited but distinct regions of synteny with *Tokpelaia* from *Tetraponera* ants.

### 2.2. Genomic features and metabolic functions in *Tokpelaia* symbionts from planthoppers and ants

The genomic contents comparisons among planthopper and ant symbionts revealed large differences among bacterial strains and host clades (Fig. 3). Generally, *Tokpelaia* strains devote a large portion of their genome contents to the metabolism of carbohydrates, amino acids, cofactors and vitamins, nucleotides, and energy metabolism, as well as to membrane transport. Interestingly, Cixiidae symbiont strains consistently encode much greater numbers of carbohydrate metabolism genes than others. In turn, the numbers of amino acid and cofactor/vitamin biosynthesis genes were relatively consistent except for the genome-reduced planthopper symbiont clade and a single ant strain (JR021), where they were substantially reduced. Interestingly, secondary metabolism genes are consistently much less abundant in planthopper symbionts than in ant symbionts.

**Fig. 3.**
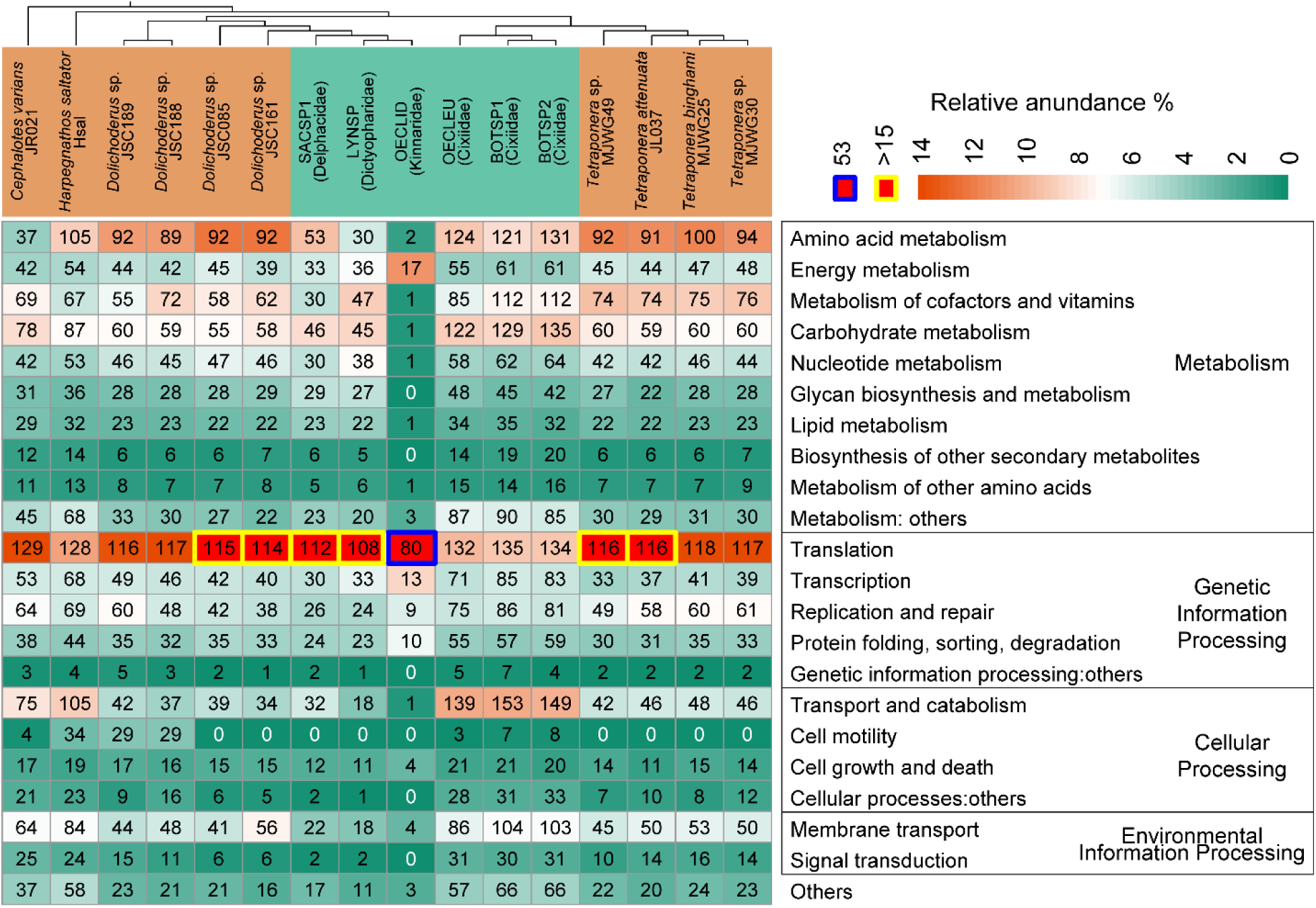
Relative abundance of KEGG pathways in *Tokpelaia* genomes. For each genome and each functional category, the numbers in the squares represent the number of genes in that category, and the background colors represent the proportion of the genes in that category relative to all annotated genes in the genome.

As the number of translation-related genes is comparable across all but the tiniest genome, their relative proportion negatively correlates with the genome size. In the tiniest *Tokpelaia*-OECLID genome, genetic information processing genes make up 61.8% of the total gene number, as despite it losing many genes in that category, much greater losses occurred in other categories. This included the elimination of all genes involved in essential amino acid biosynthesis, with only two non-essential amino acid metabolism genes (*cysE* and *cysK*) and one vitamin biosynthesis gene (*cobA*) remaining.

Comparative analysis of nutrient biosynthesis pathways across the sequenced planthopper *Tokpelaia* genomes indicated a strong similarity among symbionts of Cixiidae (BOTSP1, BOTSP2, and OECLEU) and those of herbivorous *Dolichoderus* and *Tetraponera* ants (Fig 4), whereas strains from LYNSP and SACSP1 had substantially reduced biosynthetic repertoire. Most strikingly, the highly reduced OECLID retained very little nutrient biosynthetic capacity.

**Fig. 4.**
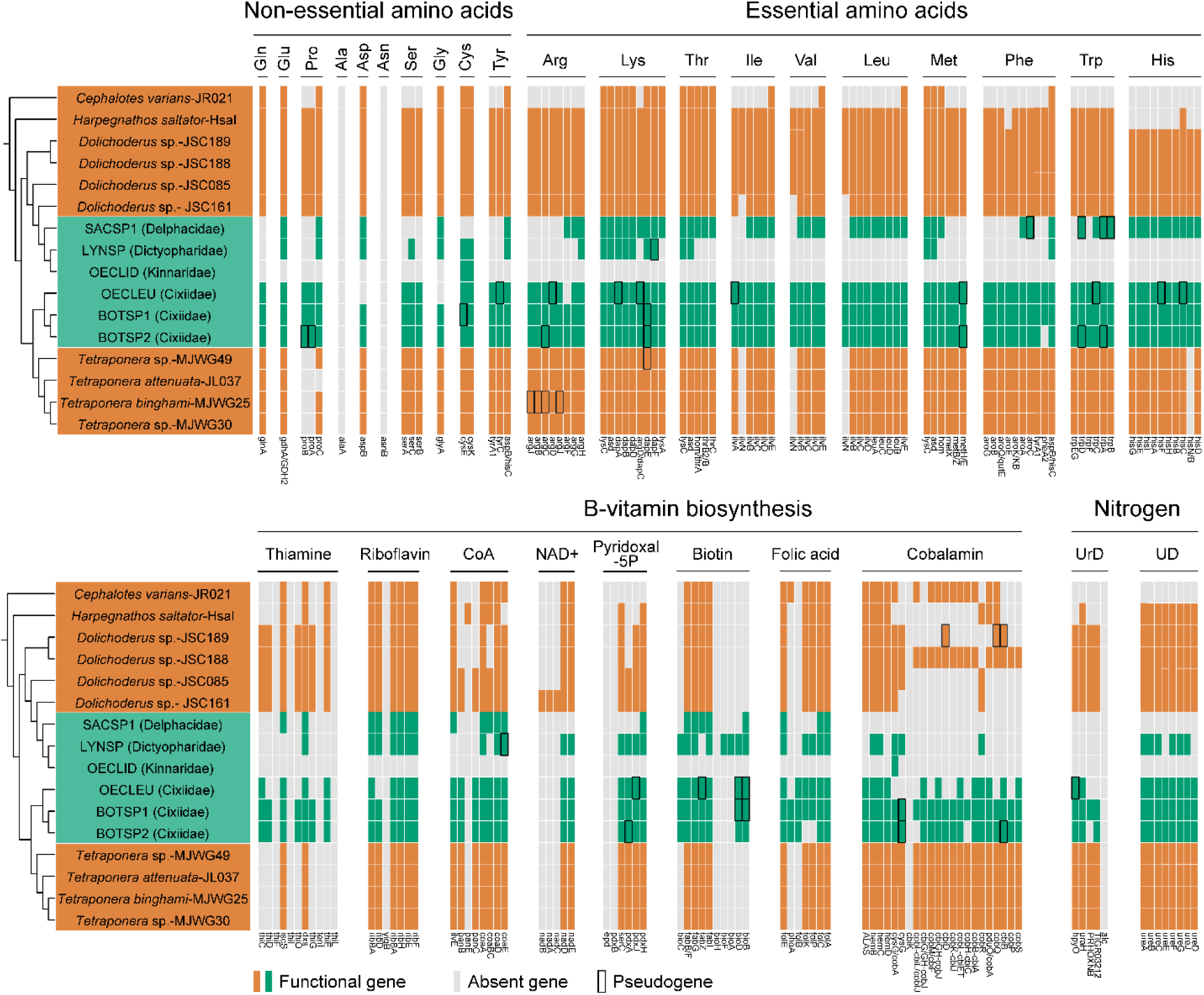
Summary of *Tokpelaia* biosynthetic functions. Genes associated with nitrogen cycling and cofactor biosynthesis in the genomes of planthopper (green) and ant (orange)-associated *Tokpelaia*. Each bar represents a single gene with an abbreviation at the bottom. Genes are classified as present (green or orange) and absent (light gray); putative pseudogenes, with disrupted open reading frames, are placed in a black frame. A cladogram based on the phylogenomic relationships of *Tokpelaia* is shown on the left, with labels in the same order as in Fig. 2A (where full host names are provided). UrD: Uric acid degradation, UD: Urea degradation.

Specifically, amino acid biosynthesis pathway comparisons revealed that the Cixiidae planthopper and *Dolichoderus* and *Tetraponera* ant symbionts retain very similar gene sets, spanning nearly complete pathways for the biosynthesis of non-essential and essential amino acids with the exception of alanine-synthesizing transaminase (*alaA*) and asparagine synthase (*asnB*). *Tokpelaia* of BOTSP2 is the only one missing prephenate dehydratase (*pheA2*), which catalyzes the decarboxylation of prephenate to phenylpyruvate in the phenylalanine pathway. The *Tokpelaia* of OECLEU lacks three additional genes: *aspB* and *glyA* for aspartate and glycine synthesis, and *argF* involved in arginine biosynthesis. Notably, however, many of the genes in *Tokpelaia* genomes of BOTSP1, BOTSP2, and OECLEU scattered across pathways, appeared pseudogenized (Fig. 4). In contrast, *Tokpelaia* of LYNSP and SACSP1 have both lost most amino acid biosynthesis genes. The LYNSP strain lacks the complete pathways for any essential amino acid, retaining six of nine genes for lysine biosynthesis and complete pathways for four non-essential amino acids: glutamate, aspartate, glycine, and serine. The SACSP1 strain retains complete pathways for three essential amino acids (lysine, histidine and threonine) and three non-essential amino acids (glutamate, aspartate, and glycine), along with partial pathways for leucine, isoleucine, and valine. This reduced biosynthetic capacity resembles strain JR021, a culturable member of conserved multi-partite microbiota of a herbivorous *Cephalotes* ant (Hu, et al. 2018). The highly reduced OECLID strain encoded only two genes involved in the biosynthesis of one non-essential amino acid, cysteine.

Besides amino acids, the *Tokpelaia* symbionts display a broad vitamin biosynthetic capability, with substantial variation among strains, and again a substantial similarity among Cixiidae, *Dolichoderus,* and *Tetraponera* symbionts. For thiamine biosynthesis, BOTSP1 and BOTSP2 symbionts encode the same set of seven genes as *Dolichoderus* strains, whereas all other strains encode a more reduced set. All *Tokpelaia* strains except OECLID symbiont retain genes for riboflavin (vitamin B2) biosynthesis, only missing *yigB* gene, with OECLEU additionally lacking *ribD*. For CoA and pyridoxal-5P, *Tokpelaia* symbionts of Cixiidae planthoppers encode similar gene sets as those of *Tetraponera* ants, with *Dolichoderus* and LYNSP-SACSP1 symbionts encoding fewer. In turn, the set of four biotin (vitamin B7) biosynthesis genes is also highly conserved among *Tokpelaia* strains (with a single gene loss in LYNSP), but all planthopper symbionts encode a set of additional genes involved in the production of this vitamin. Interestingly, BOTSP1’s *Tokpelaia* is the only one retaining the full pathway from purines to folic acid, with all other strains having experienced gene losses. Finally, cobalamin, a key coenzyme involved in the metabolism of branched-chain amino acids and methionine, has the longest synthesis pathway of 22 steps, all but one of which (*cbiK*) is encoded by *Tetraponera* and BOTSP1 symbionts. Here, BOTSP2 and one of the *Dolichoderus* strains lost more genes, and all others have a more substantially reduced set. The highly reduced OECLID symbiont only encodes one vitamin biosynthesis gene, *cobA*, in the cobalamin biosynthesis pathway.

Similarly, *Tokpelaia* from Cixiidae planthoppers and most ants (*Dolichoderus*, *Tetraponera*, and *Harpegnathos saltator*) harbor a complete urease gene cluster (ureABCDEFGJ) for urea degradation. However, the LYNSP strain lacks two accessory components (ureE and ureJ), and SACSP1 has lost the entire gene cluster, same as the *Cephalotes* ant symbiont JR021 (Fig. 4). On the other hand, while the pathway for uric acid degradation is retained in *Tokpelaia* from *Dolichoderus* and *Tetraponera* ants, BOTSP1 is the only planthopper whose symbiont retains the entire pathway, with partial or complete losses in other species (Fig. 4). Again, none of the nitrogen recycling genes were found in *Tokpelaia*-OECLID.

The close inspection of the OECLID symbiont gene set revealed, outside of genetic information processing genes, two substantial functions that were retained (Fig. S4, Tab. S7). The first is energy metabolism, including components of the electron transport chain (17 genes related to cytochrome c oxidase, ATP synthase, and electron transfer), the second is iron–sulfur cluster assembly. The only two amino acid (cysteine) biosynthesis-related genes present in this genome are also involved in iron-sulfur cluster assembly (Fig. S5).

### 2.3. The degeneration of ancient symbionts is often associated with the *Tokpelaia* acquisition

Within metagenomic assemblies of the six *Tokpelaia*-hosting planthopper species we also identified other bacteria (Fig. 1B, Tab. S2) (Michalik, et al. 2025). Specifically, within all studied insects, we identified rRNA sequences of *Vidania*, and in five of them (all except SACSP1), of *Sulcia* – two ancestral symbionts of planthoppers. BOTSP1, BOTSP2 and OECLID also contained the facultative symbiont *Wolbachia*. LYNSP carried *Rhodanobacter*, whereas SACSP1 hosted *Sodalis* as an additional symbiont (Fig. 1B). For most of these symbionts, we recovered genomic bins with acceptable completeness and contamination statistics (Tab. S2). Intriguingly, in Cixiidae species, *Vidania* and *Sulcia* genomes were fragmented and apparently incomplete, unlike in the three other *Tokpelaia*-hosting species and the vast majority of previously studied planthoppers (Michalik, et al. 2025). Further, in BOTSP2, *Wolbachia* was not successfully reconstructed through binning; we identified matching contigs via BLASTn, but because of high contamination scores for this set, as computed by CheckM (72.81%), we decided to exclude it from analyses.

The ancient nutritional endosymbionts’ genome organization and functions varied among the studied species (Fig. 5). Despite its fragmentation in Cixiidae, *Sulcia* nutritional functions were conserved in the five planthoppers that retained it, always comprising full pathways for valine, leucine, and isoleucine biosynthesis. In contrast, *Vidania* genome retained ancestral organization and the ability to produce seven essential amino acids in only two species, LYNSP and OECLID. In the third, SACSP1, *Vidania* genome was reduced to provisioning phenylalanine (Michalik, et al. 2025). In turn, in all three Cixiidae species, the unusually fragmented *Vidania* strains only retained a small and variable subset of genes scattered across biosynthetic pathways. In Cixiidae and SACSP1, the gaps in conserved essential amino acid biosynthesis pathways are mostly filled by the genes encoded by other symbionts (Fig. 5). Specifically, in Cixiidae, the largely complete biosynthetic gene sets encoded in *Tokpelaia* genomes seem to fully compensate for the *Vidania* gene losses. With a few exceptions (lysine and phenylalanine), this is also the case in SACSP1; additionally, in this species, 17 genes overlapping with those of *Tokpelaia* are encoded by *Sodalis*. In LYNSP, the conserved gene sets of *Sulcia* and *Vidania* are almost entirely duplicated by *Rhodanobacter*, with *Tokpelaia* providing further scattered functions. Finally, in OECLID, two *Wolbachia* strains encode a set of biosynthetic genes that largely overlap with those of ancient symbionts.

**Fig. 5.**
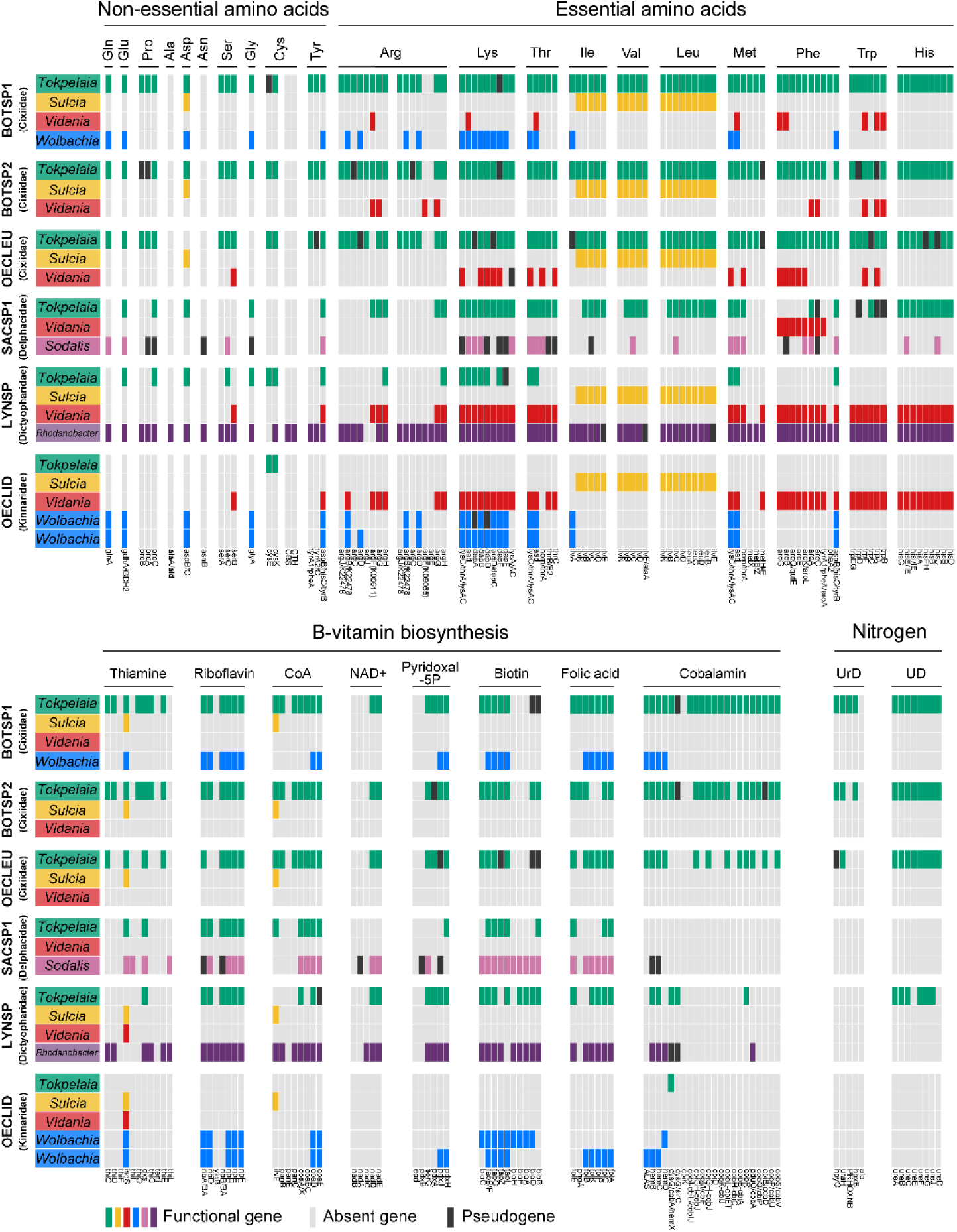
Genomic complementarity in nutritional provisioning among planthopper symbionts. Amino acid and B vitamin biosynthesis gene distribution among genomes of different symbionts that were identified in metagenomes of six *Tokpelaia*-hosting planthopper species. Genes are classified as functional genes, absent (light gray), and pseudogenes (black).

Species differed substantially in terms of B vitamin and cofactor biosynthesis gene sets and their distribution across symbiont genomes (Fig. 5). In Cixiidae species, *Tokpelaia* encodes large numbers of genes across different pathways, with *Wolbachia* duplicating these functions in BOTSP1. In SACSP1 and LYNSP, *Sodalis* and *Rhodanobacter*, respectively, encode the greatest number of genes, with *Tokpelaia* duplicating many of them. Finally, in OECLID, its two *Wolbachia* genomes encode multiple functions, and *Tokpelaia* contributes one gene. Across all these species, few vitamin biosynthetic pathways appear complete, but we note that the assembly fragmentation may have prevented the detection of some genes, and they may also be encoded within the host genomes.

## 3. Discussion

When famously writing about ‘a veritable fairyland of insect symbiosis’, Paul Buchner (1965) referred to the huge variety of bacterial symbionts of Auchenorrhyncha that accompany, and sometimes have replaced, their ancient, degraded nutritional endosymbionts. Of these, the enterobacterial genus *Sodalis* – apparently, the most broadly distributed across the taxonomic diversity of insects – may be the most widespread but also best understood in terms of genomic evolution and nutritional contributions to the host biology (McCutcheon, et al. 2019; Renoz, et al. 2023). Through our research, the family *Bartonellaceae* joins the ranks of Auchenorrhynchan co-symbionts, showing patterns of the distribution, evolution, and function not unlike *Sodalis*, but casting a new perspective on microbial symbioses of hemipterans, ants, and other insects.

### 3.1. Bartonellaceae host range broadens

*Bartonellaceae* are primarily known as pathogens of mammals, often vectored by blood-sucking insects (Engel, et al. 2011). As an example, *Bartonella bacilliformis* causes the devastating and often lethal Carrion disease in humans (Pons, et al. 2016), while *Bartonella henselae* causes persistent and usually asymptomatic infections in cats, vectored by cat fleas (Chomel, et al. 1996). The discovery of the bee gut symbiont *Bartonella apis* (Segers, et al. 2017) and the identification of more divergent strains colonizing diverse ants, classified to the *Candidatus* genus *Tokpelaia* (Russell, et al. 2009; Bisch, et al. 2018; Ma, et al. 2025), have changed the understanding of the biological spectrum of this bacterial group. Our discovery of planthopper-associated *Bartonellaceae*, fitting right within the *Tokpelaia* clade and with at least four independent original host association events, further broadens the list of confirmed hosts. The plant-based diet of many *Bartonellaceae*-hosting ants and planthoppers, but also the mutualistic interactions among many ants and hemipterans, including planthoppers (Ivens, et al. 2017; Bourgoin, et al. 2023), suggest potential avenues for parallel acquisition of related microbes by these groups, or their exchange among groups. However, we already know that *Bartonellaceae* host list is longer: for example, a divergent clade commonly colonizes stored-product mites (Xiong, et al. 2024), and *Bartonellaceae* 16S rRNA sequences were identified in a scorpion and in heteropteran bugs (Matsuura, et al. 2012; Martinez, et al. 2019). We expect that a systematic survey across diverse arthropods will reveal these bacteria in a substantially broader range of hosts. At the same time, as shown recently, systematic screens across related species and within species, microscopy, transcriptomics, and/or experiments, can clarify the nature of their associations with diverse hosts (Xiong, et al. 2024; Ma, et al. 2025).

Overall, our findings support the view of *Bartonellaceae* as relatively broadly distributed across animals, and perhaps a wider range of environments. We suspect that some strains are relatively versatile and capable of synthesizing a wide range of nutrients, which enable them to move across and colonize new host lineages and then potentially evolve into more specialized associations. Some of these, including the mammal-pathogenic *Bartonella* clade, may retain the ability to switch hosts, while others may become strictly heritable, promoting genomic reduction and the loss of functions not essential to the specific host. The latter evolutionary trajectory agrees with the emerging consensus on the evolution of some other key insect-symbiotic bacteria, as exemplified by the enterobacterial genus *Sodalis* (McCutcheon, et al. 2019). It includes strains such as *Sodalis ligni* that live in dead wood (Tlaskal, et al. 2021)*; Sodalis praecaptivus* isolated from a human wounds following impalement on a branch (Clayton, et al. 2012); and diverse strains that have independently established heritable nutritional symbiotic relationships with a wide variety of insect lineages (Renoz, et al. 2023). Such heritable associations typically lead to rampant genomic changes, increased evolutionary rates, and rapid gene losses. Some of the degenerating symbionts may reach a stable state where gene loss slows to nearly a halt – as evidenced by high similarity among genomes of *Buchnera*, *Blochmannia*, or *Baumannia*, all likely derived from *Sodalis* tens or hundreds of millions years ago (McCutcheon, et al. 2019).

### 3.2. Planthopper symbionts represent contrasting genomic evolutionary outcomes

Variation in planthopper-associated *Tokpelaia* genome size and contents indicates their different lifestyles and varying effects on the host biology, paralleling the range of associations of *Sodalis*-allied symbionts.

On the one hand, the three *Tokpelaia* strains infecting Cixiidae planthoppers have the largest genomes (∼2Mb) among the *Tokpelaia* strains studied so far (0.96-1.9Mb), retaining a broad range of amino acid and vitamin biosynthesis genes. This suggests a substantial contribution to the host nutrition, comparable to that of ant-associated *Tokpelaia* (Bisch, et al. 2018; Ma, et al. 2025). Further, these strains’ positions on short branches in the *Tokpelaia* phylogeny and high GC contents suggest proximity to the putative ancestral state of other associations with diverse ants and planthoppers. This resembles the versatile *Sodalis praecaptivus*, whose function-rich, high-GC genome is much larger than most insect-symbiotic *Sodalis* strains, and consistently falls near the base of *Sodalis* phylogenies, among variable-length branches representing insect-symbiotic strains that departed from this ancestral state (McCutcheon, et al. 2019).

On the other hand, the *Tokpelaia* genomes of LYNSP and SACSP1 planthoppers are substantially reduced (632-677 Kb), to the level comparable with *Sodalis*-derived long-term obligate nutritional symbionts of aphids (B*uchnera*, 412-671 kb), carpenter ants (*Blochmannia*, 705-792 kb), sharpshooter leafhoppers (B*aumannia*, 633-759 kb), or tsetse flies (Wigglesworthia, ∼700 kb) (Bennett, et al. 2015; Bing, et al. 2017; Chong, et al. 2019). Despite the massive variation in their age of association, tissue localization, transmission mechanisms, and biosynthetic capacity, all these gamma-proteobacterial symbionts provide deficient nutrients that make them virtually indispensable to their hosts (Feldhaar, et al. 2007; McCutcheon, et al. 2009; Bing, et al. 2017). We suspect that LYNSP and SACSP1 *Tokpelaia* strains play comparable roles in planthoppers. As both strains have lost multiple nutritional genes that are retained by other symbionts, their specific situation may particularly resemble that of *Baumannia* (Wu, et al. 2006), or *Buchnera* from *Cinara* aphids (Manzano-Marin, et al. 2016), or endobacterial symbionts of mealybugs (Husnik and McCutcheon 2016): all of them are complementary in their amino acid biosynthetic capacity to other symbionts residing within the same host insect.

In our view, the most striking discovery is the tiny, 158-kb genome of the *Tokpelaia* symbiont of the Kinnaridae planthopper *Oeclidius* sp. (OECLID), comparable in size to the most reduced known insect endosymbionts: *Sulcia*, *Vidania*, *Nasuia*, *Hodgkinia*, *Tremblaya*, or *Carsonella* (McCutcheon, et al. 2024). All these bacteria colonize sap-sucking hemipterans, where they have well-defined roles as providers of essential amino acids and sometimes vitamins. This does not appear to be the case with *Tokpelaia*-OECLID – even if, as shown in a mealybug, one nutrition-related biosynthetic gene (*cobA* in the current case) might potentially be enough to justify long-term symbiosis (Garber, et al. 2024). But in fact, OECLID symbiont’s genetic repertoire resembles much more closely the endosymbionts of certain protists. Specifically, the 158-kb genome of a Verrucomicrobium, *Ca.* Organicella extenuata, symbiont of an *Euplotes* protist inhabiting a hypersaline Antarctic lake, also retains the capacity for genetic information processing but not for the biosynthesis of amino acids or vitamins (Williams, et al. 2021), with fatty acid synthesis and iron-sulfur (Fe-S) cluster assembly identified as its likely contribution to the host biology. Fe-S cluster assembly has not been identified as critical to sap-sucking insects’ biology. Thus, the nature of the association, significance, and evolutionary history of this unusual planthopper-associated bacterium that conflicts with the current understanding of insect heritable endosymbioses remains a mystery and, unfortunately, it cannot be resolved based on a single mid-coverage metagenome alone. To further complicate conclusions, from OECLID metagenome, we also reconstructed a low-coverage rRNA sequence representing an Entomophthorales fungus with 96.6% identity to entomopathogens *Pandora delphacis* and *Furia pieris*, and we cannot exclude the possibility that *Tokpelaia*-OECLID actually associates with the planthopper’s pathogen (Matsuura, et al. 2018; Siehl, et al. 2024).

### 3.3. *Bartonellaceae* significance in the biology of planthoppers and their other symbionts

We know that heritable nutritional endosymbionts in diverse insect systems produce essential nutrients deficient in their hosts’ diets, thus being indispensable to the hosts. At the same time, they drive the evolution of hosts’ dependence on symbiont nutritional contributions, even as these symbionts degrade and loose efficiency – a process aptly described as ‘the evolutionary rabbit hole’ (Bennett and Moran 2015). Further, they drive the complementary loss of functions by their co-symbionts (McCutcheon, et al. 2009; Husnik and McCutcheon 2016), forcing the hosts to manage an increasingly specialized symbiotic consortium. We think that these general statements accurately describe the biology and significance of most planthopper-associated *Tokpelaia* strains that we have identified, which in all cases, belong to multi-partite symbiotic consortia with complementary biosynthetic capabilities (Michalik, et al. 2021). Specifically, in a delphacid SACSP1, the *Bartonellaceae* symbiont largely fills up the nutritional gap resulting from *Sulcia* loss and *Vidania* genome reduction to the smallest-known state, while in Cixiidae, they seem to compensate for the apparent degeneration of *Vidania.* While such complementarity cannot be seen in LYNSP, and in OECLID it is limited to a single gene, it is tempting to think of *Tokpelaia* as the common driver of the functional loss by ancient symbioses. At the same time, other symbionts are likely to be driving the gene loss in *Tokpelaia.* We propose that through alternative evolutionary routes, symbiotic complexes involving *Tokpelaia* have reached, or are on a path towards, complementarity in the biosynthesis of largely the same set of nutrients as most other planthoppers. Depending on the order of arrival and the genomic properties of bacteria that were in planthoppers before, the independently acquired *Bartonellaceae* have assumed different positions and roles in the symbiotic consortium.

We are not aware of any major ecological transitions in planthopper lineages represented by *Tokpelaia*-infected species. Such changes, exemplified by Typhlocybinae leafhoppers that switched to feeding on parenchymal cell cytoplasm (Buchner 1965; Kobiałka, et al. 2025) and in Derbidae and Achilidae planthoppers that switched to feeding on fungi during nymphal stages (Michalik, et al. 2025), seem to have enabled major changes and functional losses in their microbiota.

However, in all presented systems, achieving the full understanding of host-microbe associations would require a much broader set of data than what we had available - a combination of within-species microbiome surveys, genomic comparisons among closely related species, microscopy, and, ideally, experiments. Unfortunately, these approaches are not easy to implement on unidentified non-model species sampled on different continents years ago, which was the case here. Nevertheless, this study demonstrates that a broad genomics survey alone can help discover new biological states and processes that change our understanding of biology and evolution of significant clades of organisms. In the age of genomics, such broad surveys should become an important tool for the discovery and characterization of biodiversity and biological interactions that shape it.

## 4. Methods

### 4.1. Study material and data

For the study, we used a set of metagenomic datasets for 149 taxonomically diverse planthoppers, presented in our previous work that focused on phylogenomic relationships of these planthoppers and comparative genomics of their *Sulcia* and *Vidania* symbionts (Deng, et al. 2025; Michalik, et al. 2025). Briefly, insects were wild-caught from their natural habitats around the world between 1996 and 2022. Following identifications by taxonomic experts, the DNA extracted from dissected abdominal tissue of single individuals, or in a few cases, dissected bacteriome tissue, was used for metagenomic library preparation and sequencing on Illumina platforms. Collection metadata, reagents, and sequencing platforms for the entire metagenomic collection are provided in the abovementioned publications. For the six specimens within which we detected *Tokpelaia*, listed in Table S1, abdominal tissue was used for DNA extraction using the Bio-Trace DNA Purification Kit (Eurx, Poland), and sequencing libraries were prepared and sequenced by Novogene Ltd. on NovaSeq 6000 platform using a 300-cycle kit.

### 4.2. Bioinformatics analysis

To detect *Bartonellaceae* within the sequenced metagenomes, we systematically examined 16S rRNA sequences assembled using phyloFlash v3.4.2 (Gruber-Vodicka, et al. 2020), validating all those without close matches to recognized bacteria through phylogenetics and blastn comparisons against NCBI databases. In all cases where these data suggested Hyphomicrobiales infections, we followed with the analysis of metagenomic assemblies obtained using Megahit v1.2.9 (maximum k-mer size = 255, min contig size= 1000) (Li, et al. 2015). To obtain the draft genomes of *Bartonellaceae* and other bacteria present in these samples, we binned the metagenomic assembly data using the metaWRAP pipeline (Uritskiy, et al. 2018), incorporating three binning tools: CONCOCT, MaxBin, and metaBAT. The resulting bins were refined with metaWRAP’s “Bin_refinement” module, applying a minimum completeness threshold of 70% and a maximum contamination limit of 10%. To avoid the omission of genomic contigs, assemblies were further examined using BLASTn and BLASTx searches against the NCBI database, and separately against published and unpublished genomes of different insect-symbiotic bacteria. Only contigs consistent with the bin classification, and meeting criteria for read coverage and GC content were retained (Tables S2, S3). Bin completeness and contamination were evaluated using CheckM v1.0.12 (Parks, et al. 2015), and taxonomic classification was performed using the Genome Taxonomy Database via GTDB-Tk v2.1.122 (Chaumeil, et al. 2019). Average amino acid identity (AAI) was calculated using CompareM v0.1.2 (https://github.com/dparks1134/CompareM).

Contigs of bacteria other than *Sulcia* and *Vidania* were annotated with Prokka v1.14.6 (Seemann 2014), and gene functions were assigned to KEGG Orthology (KO) pathways using KofamScan (Aramaki, et al. 2020) and the online tool eggNOG-mapper (Tab. S6) (Cantalapiedra, et al. 2021). Based on these annotations, metabolic pathways of the symbiotic bacteria were manually reconstructed using the KEGG database. Genomes were visualized using Proksee (Grant, et al. 2023). Genome synteny analysis was performed using Promer tool for Mummer v3.07, with default settings (Kurtz, et al. 2004). Pseudofinder v1.1.0 (Syberg-Olsen, et al. 2022) software was used to predict pseudogenes in the genomes of symbiotic bacteria. In turn, all confirmed *Sulcia* and *Vidania* contigs and genomes were annotated using a custom pipeline (Łukasik, et al. 2018) adopted for planthopper symbionts recently (Michalik, et al. 2025). The pipeline first extracted all Open Reading Frames (ORFs) and translated them into amino acid sequences. These ORFs were then searched using HMMER v3.3.1 (Eddy 2011) (hmmer.org) against a custom database containing manually curated protein-coding and rRNA genes from previously characterized *Sulcia* and *Vidania* genomes. Ribosomal RNA genes were detected with nhmmer implemented in HMMER v3.3.1, and tRNA genes were identified with tRNAscan-SE v2.0.7 (Chan, et al. 2021) Based on the relative length to the reference genes, protein-coding genes were classified as functional (>90%), putative pseudogenes (>60%), or pseudogenes (<60%). Any ORFs longer than 300 bp with no significant similarity to any reference genes were further examined by querying UniProt (The UniProt 2021) and NCBI databases and compared carefully to the top hits. Genes lacking any annotations were marked as “hypothetical”.

### 4.3. Phylogenetic analyses

To investigate the evolutionary origin of *Bartonellaceae* bacteria associated with planthoppers, we compared phyloFlash-reconstructed 16S rRNA sequences against top hits of insect and environmental origin, especially published ant-associated *Tokpelaia* sequences. All sequences were aligned using MAFFT v7.526 (Rozewicki, et al. 2019), with divergent regions removed using Gblocks v0.91b (Talavera and Castresana 2007). Phylogenetic analysis was performed using RAxML v8.2.12 (Stamatakis 2014) under the GTRGAMMA model, with *Brucella pseudintermedia* (accession number: CP099968) designated as the outgroup. The analysis included 1000 bootstrap replicates and was visualized using iTOL (Letunic and Bork 2021).

To clarify the evolutionary relationships among the newly identified strains and other *Bartonellaceae*, we conducted a genome-wide phylogenetic analysis that included all currently published *Tokpelaia* or *Bartonella* genomes representing the major recognized clades. The proteins were clustered into families using OrthoFinder v2.5.5 (Emms and Kelly 2019), resulting in a data set of 68 single-copy orthologs. And then, proteins from each family were aligned using MAFFT v7.526 (Rozewicki, et al. 2019), with divergent and ambiguously aligned blocks trimmed using Gblocks v0.91b (Talavera and Castresana 2007). We concatenated the resulting alignments. Prottest 3.4.2 (Guindon and Gascuel 2003; Darriba, et al. 2011) was used to select the optimal amino acid substitution model, JTT+I+G+F, according to AIC and BIC criterions. Phylogenetic analysis was performed with RAxML v8.2.12 (Stamatakis 2014) using the best model and 1000 bootstrap replicates, and results were visualized and annotated using iTOL (Letunic and Bork 2021). All images were adjusted using Adobe Illustrator CC 2019.

## 5. Data availability

All raw data and genomic assemblies used in comparisons have been uploaded to NCBI databases. BioProject and BioSample accession numbers are listed in Tables S1-S2. Supplementary Tables were deposited in the public Figshare repository under the DOI: 10.6084/m9.figshare.30010978.

## 6. Acknowledgements

This study was supported by the Polish National Science Centre grant 2018/30/E/NZ8/00880 to P.Ł. and the China Scholarship Council grant no. 202406040164 to M.M.

## 7. Author Contributions

M.M., A.M., and P.Ł. conceived the study. J.D. and A.M. provided bioinformatics support and additional *Vidania* and *Sulcia* genome annotations. M.M., A.M., J.D., Y.H., and P.Ł. analyzed the data and wrote the manuscript. All authors reviewed and agreed on the manuscript.

