## Supplementary Figures for "Contrasting genomic trajectories of *Bartonellaceae* symbionts of planthoppers"

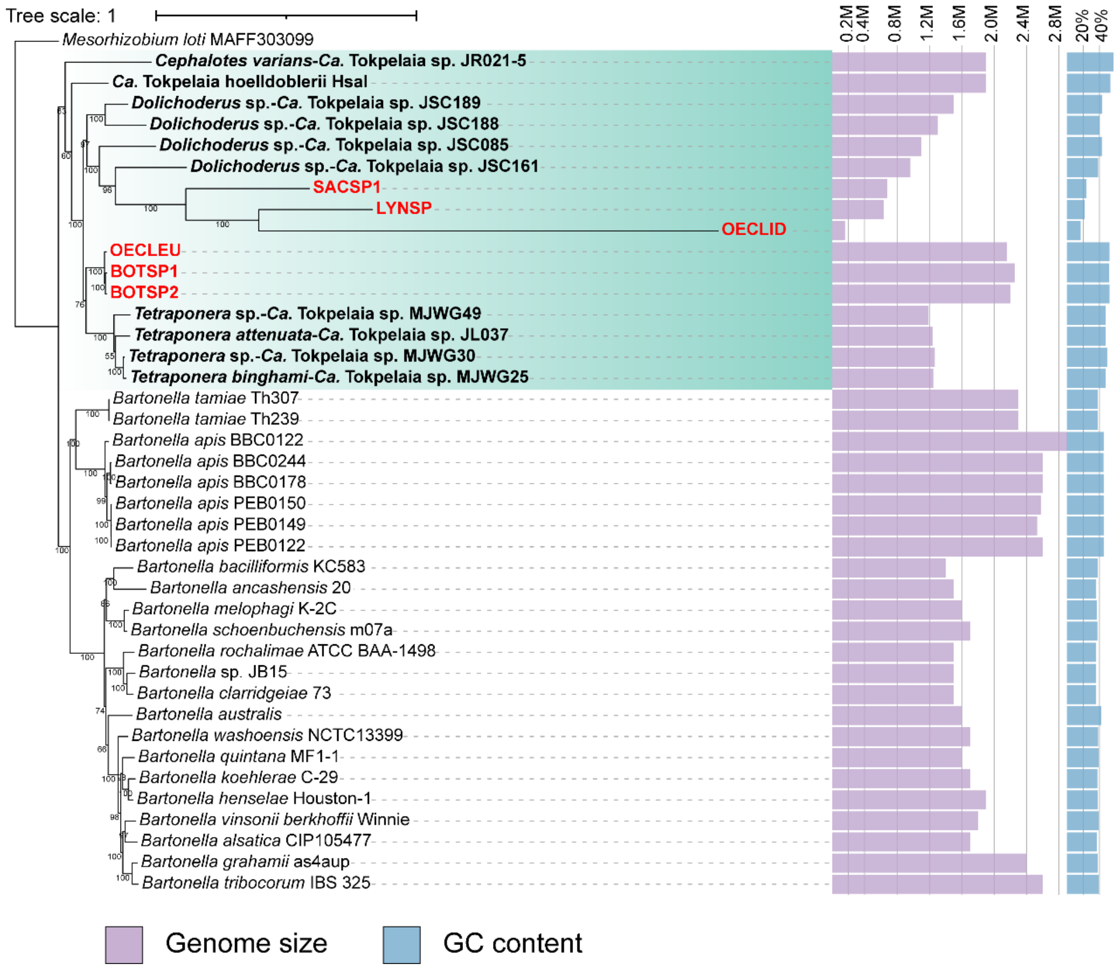


**Figure S1. Phylogenetic relationships of *Bartonellaceae* symbionts of planthoppers, ants and other closely related bacteria.** Maximum likelihood analysis of *Bartonellaceae* strains with sequenced genomes, based on 68 homologous single-copy genes. The purple bars represent the genome size, and the blue bars - the GC content. The *Tokpelaia* clade is indicated with green boxes; bold black labels indicate ant-associated, and bold red – planthopper-associated strains. Bootstrap support values above 50% are shown by the respective nodes. Unlike in Fig. 2A, OECLID branch has not been truncated.

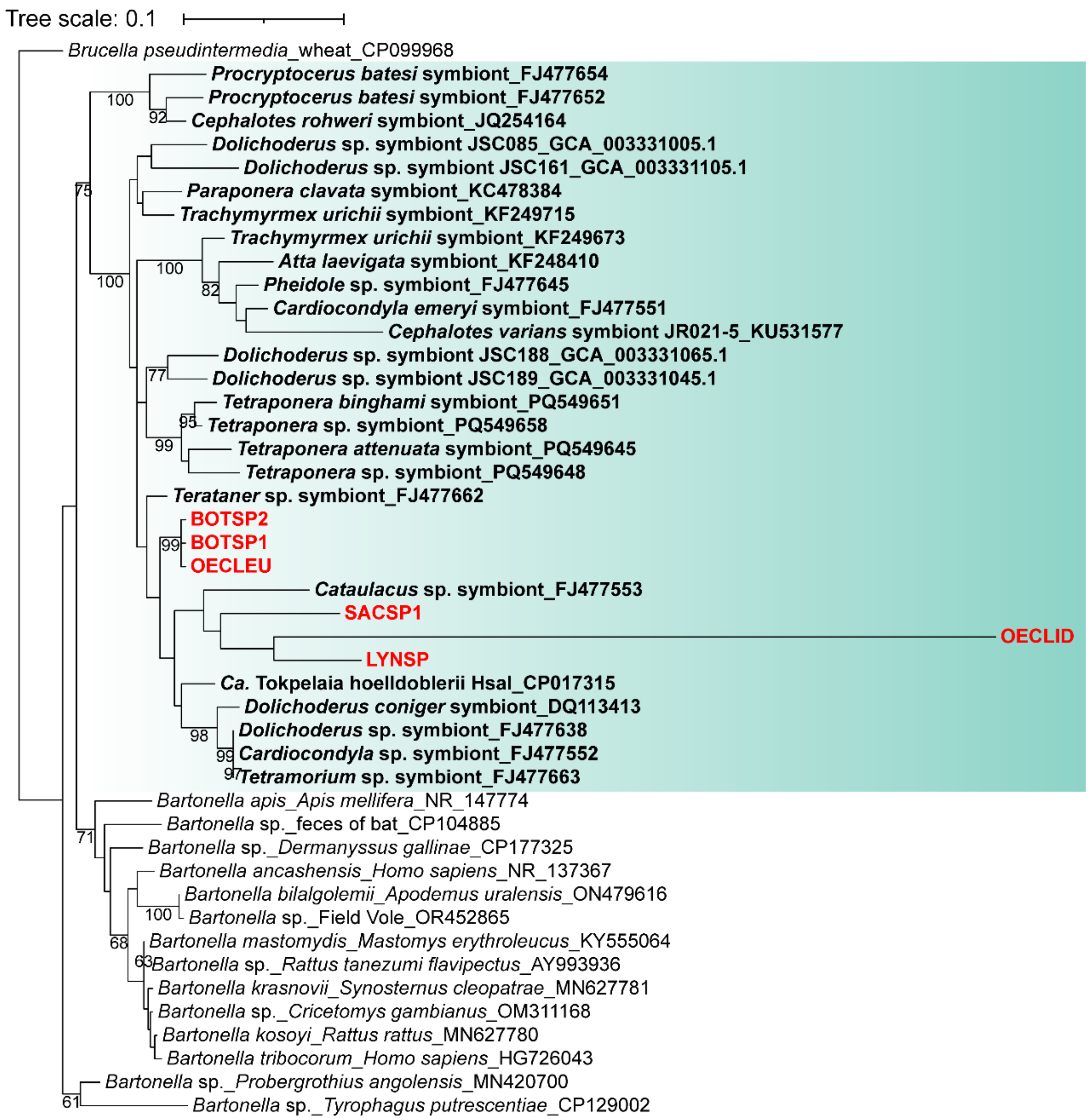


**Figure S2. Maximum likelihood phylogeny of full-length 16S rRNA gene sequences of *Bartonellaceae* associated with planthoppers, ants, and selected other hosts, derived from metagenomes and the NCBI database**. The *Tokpelaia* clade is indicated with green boxes; bold black labels indicate ant-associated, and bold red – planthopper-associated strains. Bootstrap support values above 50% are shown by the respective nodes. Unlike in fig. 2B, OECLID branch has not been truncated.

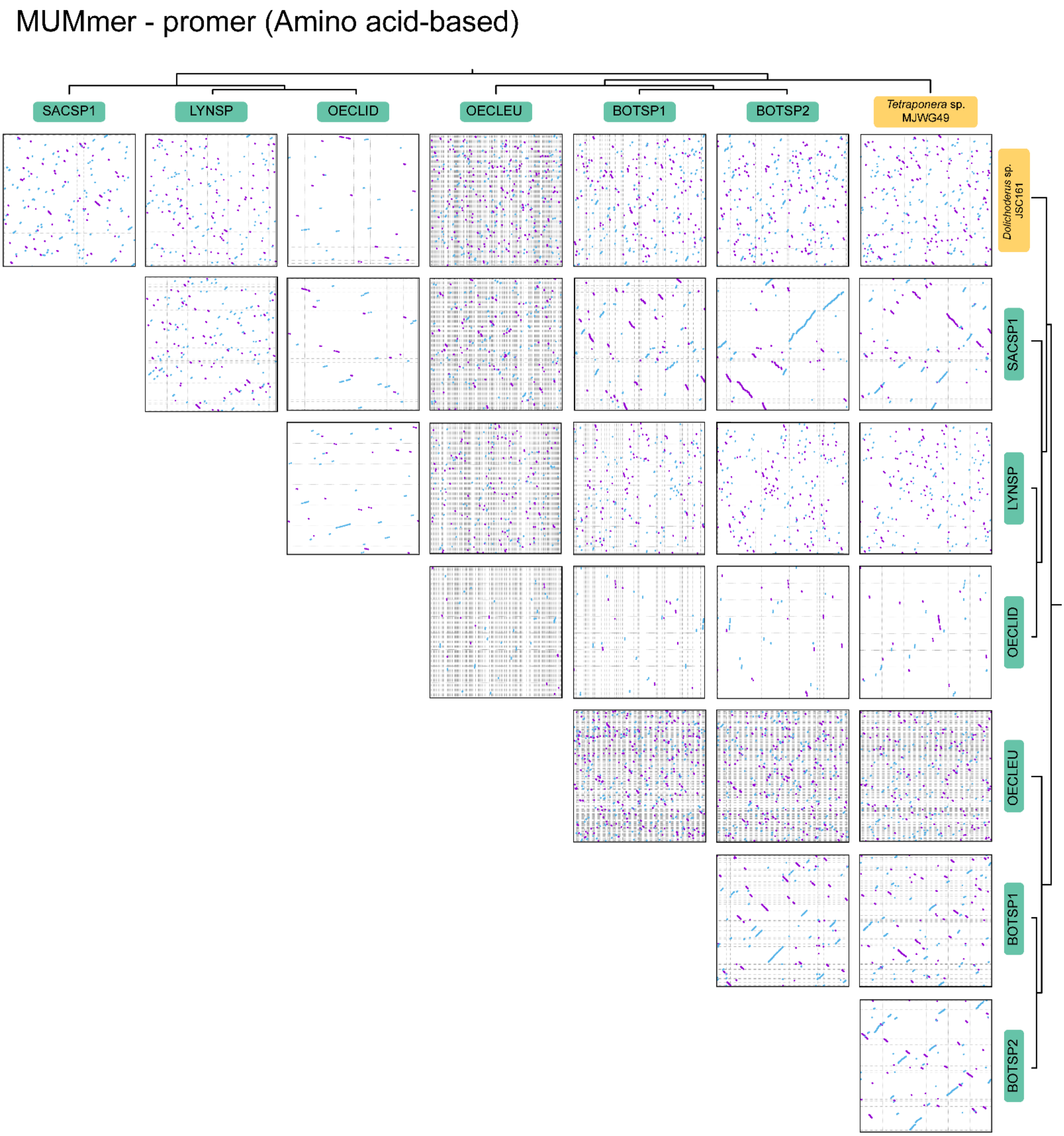


**Figure S3. Dot plots of protein-based alignments among different *Tokpelaia* strains.** Comparison of gene order in the genome contigs of *Tokpelaia* symbionts associated with planthoppers, and the genomes of *Dolichoderus* and *Tetraponera* ant symbionts. Comparisons have been done using promer v3.07, with default settings; lines and dots represent regions with significant amino acid sequence similarity, with forward matches colored in purple and reverse matches in blue.

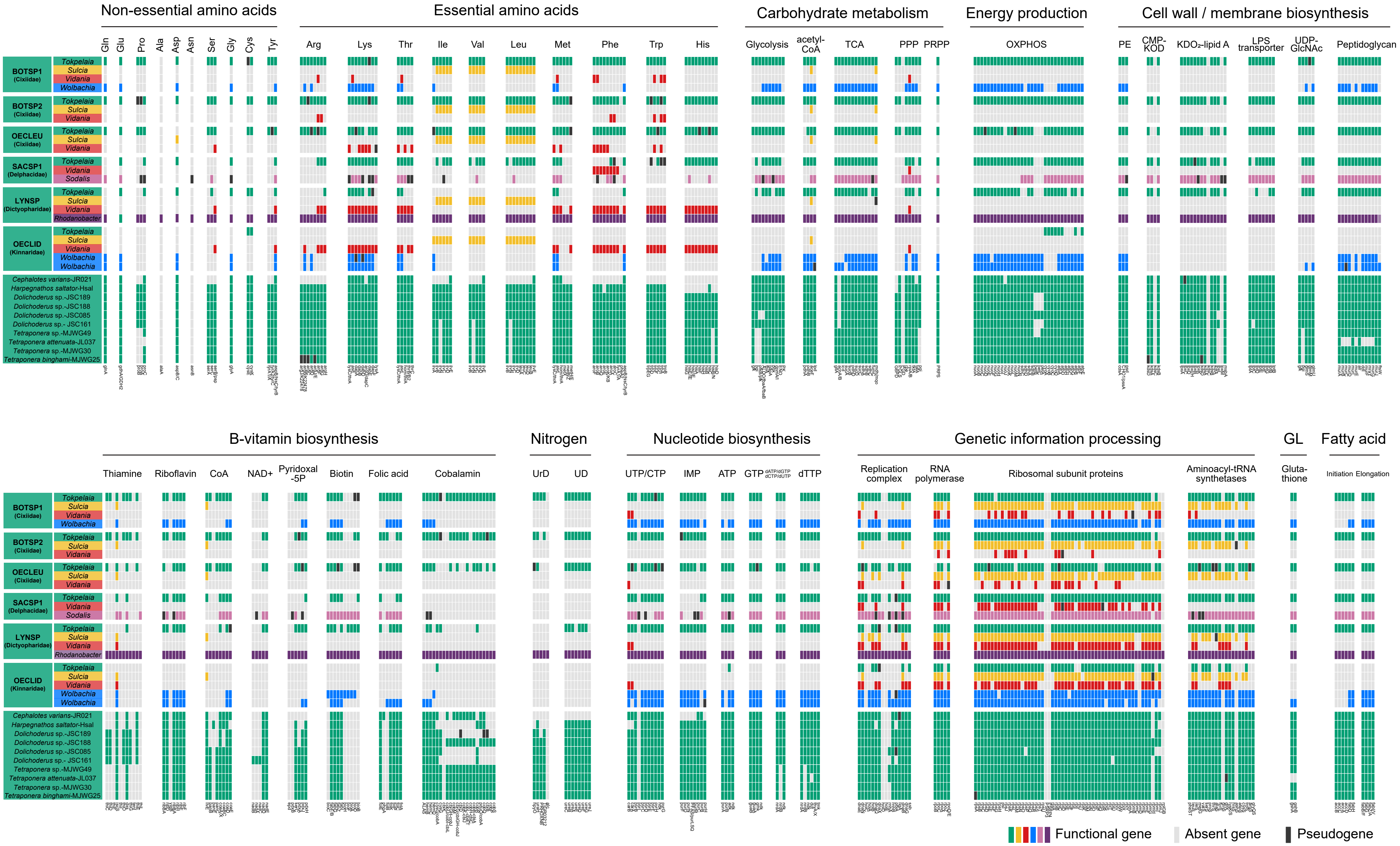


**Figure S4. The comparison of gene sets related to amino acids, vitamins, cell membrane components, nucleotide, fatty acid biosynthesis, carbohydrate metabolism, nitrogen cycling, energy production, and genetic information from bins and circular genome of symbionts of six planthopper and *Tokpelaia* symbionts of ants.** Each bar represents a single gene with an abbreviation at the bottom. Genes are classified as functional genes, absent (light gray), and pseudogenes (black).

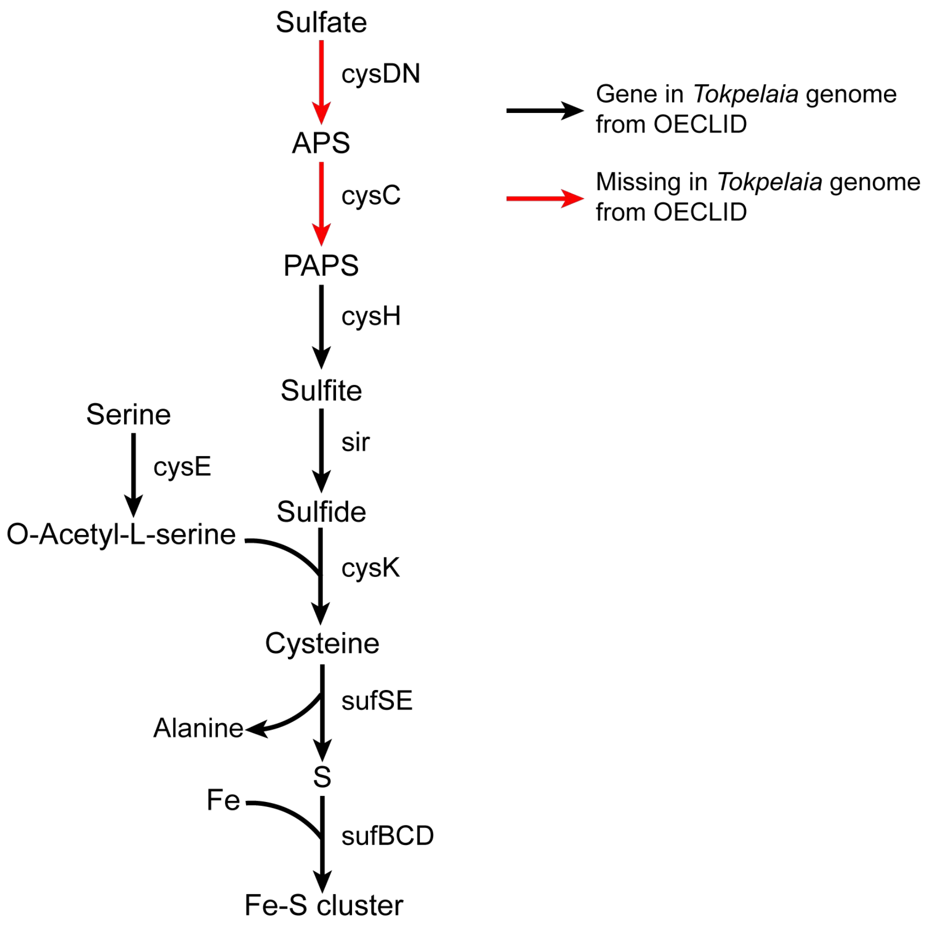


**Figure S5. Sulfur-related metabolic pathways across *Tokpelaia* symbionts of OECLID.** Black and red arrows indicate genes present and absent in the *Tokpelaia* genome of OECLID, respectively.
